# The Effects of Mindfulness on Brain Network Dynamics Following an Acute Stressor in a Population of Moderate to Heavy Drinkers

**DOI:** 10.1101/2025.09.15.676300

**Authors:** Shannon M. O’Donnell, W. Jack Rejeski, Mohammadreza Khodaei, Robert G. Lyday, Jonathan H. Burdette, Paul J. Laurienti, Heather M. Shappell

## Abstract

Previous research has found that mindfulness-based techniques are beneficial for reducing stress in heavy drinking individuals. However, the underlying neurobiology of these stress-reducing effects are unclear. Moreover, much of the research examining neurobiological correlates of mindfulness have used static functional connectivity, suggesting brain activity goes unchanged for the entire length of an MRI scan. In the current study, we used a state-based dynamic functional connectivity model to examine brain states during either a 10-minute mindfulness session or resting control that followed an individually tailored stress imagery task. Using a Hidden Semi-Markov Model (HSMM), six brain states and the associated dynamics of state traversal were estimated for the population. Participants that experienced the mindfulness session had more transitions and longer time spent in states in which the salience network was more active. Participants assigned to the control group had more transitions and increased time spent in states in which nodes of the default mode network were more active. Moreover, for control participants, increased occupancy time to SN-dominant states were associated with lower perceived stress. Using HSMM provided unique insight into network connectivity during mindful states; we believe it offers a novel approach to testing and optimizing the content of mindful-based therapies.

## I. Introduction

According to the 2023 National Survey of Drug Use and Health (NSDUH), approximately 16 million adults aged 18 and older reported heavy alcohol consumption in the past month (Substance Abuse and Mental Health Services Administration [SAMHSA], 2023). Heavy drinking is defined by the National Institute on Alcohol Abuse and Alcoholism (NIAAA) as five or more drinks on any day or ≥15 drinks per week for men and four or more drinks on any day or ≥8 per week for women (NIAAA, 2025). It is well known that stress is a key factor in alcohol consumption (Keyes et al. 2012; Sinha, 2007), as it increases the amount of alcohol consumed as well as the risk of relapse in early periods of abstinence. Moreover, acute stress has been shown to contribute to the progression of drinking behaviors, from heavy drinking to alcohol abuse to alcohol dependence (Lijffijt et al. 2014).

Two subnetworks of interest that are implicated in the pathophysiology of heavy drinking are the default mode network (DMN) and salience network (SN). The DMN includes regions such as the ventral medial prefrontal cortex (vmPFC), posterior cingulate cortex (PCC), and parts of the medial and lateral temporal cortex (Fang et al. 2021). Evidence has suggested that the DMN is predominantly active at rest and during self-referential thought and introspection (Menon, 2023). Altered activity and/or connectivity between the DMN and other cortical regions implicated in emotional regulation has been associated with increased stress-induced drinking (Zhang & Volkow, 2019). The SN has two functional hubs: the insula and the dorsal anterior cingulate cortex (dACC) which are important in allocating attentional resources to the most salient stimuli (Schimmelpfennig et al. 2023). The insula is a central component of the larger systems that process drug cues, stress, and reward (Naqvi et al. 2014; Zilverstand et al. 2018). Additionally, increased insular activity as a result of stress is associated with higher alcohol craving and consumption (Bach et al. 2024; Garavan, 2010).

Emerging research has suggested that building resilience to stress could function as a protective factor against risky drinking behaviors (Cusack et al. 2023; Kabat-Zinn, 2005). In particular, following the encounter with an acute stressor, a viable action plan could be to invoke brief mindfulness-based stress reduction (MBSR) techniques (A. Niazi & S. Niazi, 2011). The concept of mindfulness is defined as purposefully and non-judgmentally paying attention in a particular way to the present moment (Kabat-Zinn, 1994). It is suggested that mindfulness techniques increase resilience to stress by enhancing cognitive awareness which in turn leads to enhanced monitoring and processing of emotional responses (Garland et al. 2009). Mindfulness practices have been used in a variety of settings, most notably to promote recovery from stress-induced alcohol seeking and relapse prevention (Goldberg et al. 2021; Kamboj et al. 2017; Li et al. 2017; Priddy et al. 2018; Zgierska et al. 2008). The neurobiology underlying mindfulness-based stress reduction has uncovered two important subnetworks, the SN and the DMN. During meditation, decreased activity of the DMN is associated with decreased mind-wandering and rumination, both of which contribute to anxiety and depression (Calderone et al. 2024). The SN is thought to play a key role in supporting mindfulness by directing attentional resources towards the present moment and suppressing any mind-wandering from the DMN (Mooneyham et al. 2016).

However, much of the literature supporting the role of the DMN and SN in mindfulness is based on a paradigm of “static” connectivity. Static functional connectivity is calculated using correlations between blood oxygenation level dependent (BOLD) signals (Ogawa et al., 1990) over the length of the entire MRI scan (Park et al. 2017). This type of analysis assumes that relationships between brain regions remain constant throughout a scan, which can range anywhere from 5-30 minutes. This may not be the case, especially during mindfulness, as participants can shift between various states of focus was well as distraction throughout a single session. Therefore, using models that assess dynamic functional connectivity provides a unique opportunity to investigate the changes in brain connectivity throughout a mindfulness session, avoiding masking of important results due to averaging data across a session. More specifically, using a model that estimates brain “states,” represented by unique brain networks of distinct activity and functional connectivity, has proved successful in identifying neurobiology associated with alcohol-related behaviors (McIntyre et al. 2024).

Following an acute laboratory stressor, the present study aimed to investigate dynamics of the DMN and SN during an active mindfulness session versus passive resting control to identify differences in state-based characteristics and explore possible correlates with self-reported stress. The remainder of this paper focuses on the effects of mindfulness to reduce stress reactivity among individuals who engage in moderate to heavy alcohol consumption. Following an acute stressor, we hypothesized that, compared to control (a brief passive recovery session), participants randomized to a brief guided mindfulness session (active recovery group) would have higher occupancy time and transition frequency groups to states in which nodes of the SN are more active; conversely, we hypothesized that those in the control group would have higher occupancy time and transition frequency to states in which nodes of the DMN are more active. Finally, we hypothesized that longer time spent and increased transitions to SN dominant states would be associated with lower perceived stress following recovery.

## II. Materials and Methods

### i. Participants

Thirty-four moderate-to-heavy drinking individuals were recruited from the local community using a variety of advertisement techniques, such as flyers, mailers, and internet postings. The final sample consisted of 32 participants, as two participants were removed. One was removed due to having stress scores far outside the range (more than double the mean of everyone else), causing them to be highly influential data points (i.e., they had a large influence on the association between stress score and brain state occupancy time). The other was removed as they had a high number of transitions early on in each of their scans, most likely due to motion, which caused them to have a large influence on group differences for occupancy time, transition frequency, and dwell time. For completeness, results that include this participant are included in the Supplementary Material. A priori analyses determined no significant differences in the stress scores based on assigned drinking state following the recovery period (p=0.252). As such, both the normal and abstained drinking state scans were included in subsequent analyses to increase statistical power.

To be eligible for the study, female participants had to consume 1-3 drinks per day or 2-4 drinks per day for males and demonstrate these drinking behaviors for at least the past three years. During a preliminary screening, drinking behavior was confirmed using responses to the first three questions of the standardized Alcohol Use and Disorders Identification Test (AUDIT) (Saunders et al. 1993) and alcohol diary responses which assessed, more accurately, alcohol consumption patterns reported by the participants. Specifically, participants were asked to report alcohol intake (amount and time of day) for each day of one full week. Exclusion criteria included the following: current or previous diagnosis of AUD and/or psychiatric disorders, current neurological disorders, smoking > 30 cigarettes per day, consuming ≥ 500mg of caffeine per day, and positive urine drug screening (methamphetamine, cocaine, marijuana, amphetamines, opiates, and benzodiazepines). Because of the association between body mass index (BMI) and blood-alcohol concentration (BAC), BMI was restricted to a range of 18.5-35 (Wang et al. 1992). Participants also had to be right-handed, not claustrophobic, and have no contraindications to MRI.

### ii. Stress Imagery and Mindfulness

To develop scripts for the acute stress imagery session, participants completed an interview session to gather individualized descriptions of stressful stimuli. After these descriptions were collected, participants rated them using a 10-point Likert scale in which 0 = not at all stressful and 10 = the most stress they have felt in their life. Stimuli rated ≥8 were used for the imagery script development. The information obtained to create the imagery scripts was then used to create an audio recording for both of the MRI visits. Imagery scripts have been standardized and used previously as an effective technique for modeling stress in the laboratory (Sinha 2009; Sinha et al. 2003).

Following the screening visit, participants were randomized to one of two 10-min protocols: a period of active mindfulness or a passive control period which occurred immediately following the stressful imagery session in the scanner. The active mindfulness session involved subjects listening to a guided experience in mindfulness-based meditation using a script created by a co-investigator. The script focused on breath counting, accepting any negative feelings that may arise, and body centering. The passive control session involved subjects being instructed to lie quietly and rest.

### iii. MRI Study Visits

Each participant completed three visits: a baseline screening visit and two MRI visits. Following the initial phone screening, the baseline visit was conducted to obtain informed consent, verify negative drug and alcohol tests, collect self-report questionnaire data, and have participants complete a modified Timeline Follow Back (TLFB) (L. Sobell & M. Sobell, 1992). In addition, script development based on methods developed by Sinha (2009) for neutral and stressful imagery tasks were completed during this visit. Stressful scripts were developed using participants’ own descriptions of a recent event that was a “most stressful” experience. Each participant completed two scan sessions: one MRI visit in a normal drinking state, defined as typical alcohol consumption for the past 3 days, and an MRI visit in an abstained drinking state, in which a 3-day period of abstinence was imposed. Participants were randomly assigned to complete the abstained state scan first and the other half were assigned to complete the normal state first. To ensure compliance with the protocol, phone calls were made to participants before imaging visits to confirm they were adhering to the assigned drinking state. Additionally, self-report questionnaires were completed at the beginning of each imaging visit.

### iv. Scan Session Protocol

The MRI scan protocol consisted of one high resolution anatomical sequence followed by a series of functional scans with a 10-point Likert scale response in between scans to assess stress levels, imagery vividness, and alcohol craving. After the anatomical sequence, a 5-minute resting state functional scan was performed. Next, a functional scan was performed while participants listened to the neutral image script. Immediately following this scan, participants were asked to rank their stress levels and the vividness of the imagery. Next, another functional scan was collected while participants were listening to their individualized stressful imagery scripts. Again, immediately following this scan, participants were asked to rank their stress levels, imagery vividness, and alcohol craving on a scale from 1-10. Next, participants completed either a 10-minute guided mindfulness experience or were asked to lie quietly and rest for 10 minutes. Once more, participants were asked to rank their stress and alcohol craving on a scale of 1-10 immediately following the active/passive recovery sessions. These self-reported stress scores were used to examine associations between state-based characteristics during mindfulness and stress levels directly following the recovery period. Finally, another 5-minute resting state functional scan was performed. The full scanner protocol was completed twice for each participant, once in a normal drinking state and once in a period of imposed abstinence. For a visual representation of the functional portion of the scanning protocol followed in this study, see Figure 1.

**Figure 1.**
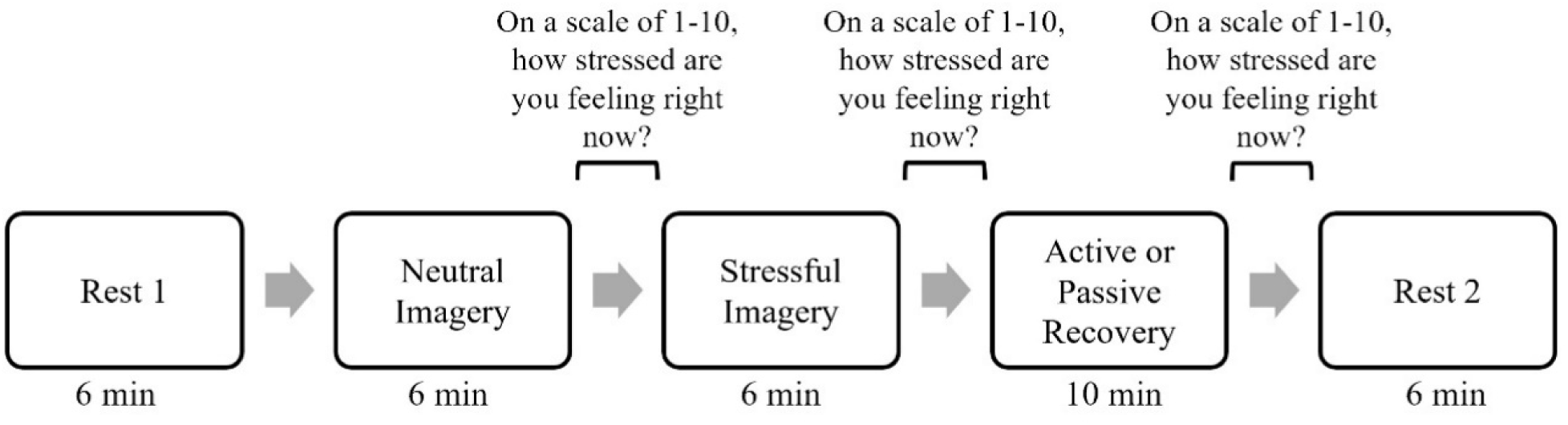
Visual representation of the functional imaging protocol used during the MRI scanning session. This full protocol was completed for each of the participants assigned drinking states.

### v. MRI Data Acquisition and Processing

Functional and anatomical imaging data was acquired on a 3T Siemens Skyra Scanner with a 32-channel head coil, a rear projection screen, and MRI compatible headphones. The imaging protocol involved a T1-weighted structural scan followed by a series of BOLD-weighted scans with varying lengths (Figure 1). For the current manuscript, we focused specifically on the active and passive recovery scans that followed the stress imagery. The high-resolution (1mm isotropic) T1-weighted anatomical sequence was performed using a single-shot 3D MPRAGE GRAPPA2 sequence with a repetition time (TR) of 2.3s, echo time (TE) of 2.99ms and 192 slices. The BOLD-weighted image sequences were acquired using an echo-planar imaging sequence with a 3.5mm x 3.5 mm x 5 mm resolution, a TR of 2s, a TE of 25ms, a flip angle of 75°, 35 slices per volume. The recovery scans used here had 297 volumes.

Data was preprocessed using Statistical Parametric mapping (SPM12) (Friston, 1994). Each participant’s T1 was coregistered to the Montreal Neurological Institute (MNI) 152 brain template (Maintz & Viergever, 1998) to enhance segmentation and warping. Unified Segmentation of the T1 images provided warping parameters to the standard space MNI template as well as brain segment images of grey matter, white matter, and cerebrospinal fluid. For each functional scan, the first 10 image volumes were removed to allow the fMRI signal to reach a steady state. Then, slice timing correction and realignment were performed. Finally, the functional scans were coregistered with the subject’s T1 image and normalized to the MNI template using the previously mentioned warp. To remove low-frequency drift and high-frequency physiological noise, a band pass filter of 0.009-0.08Hz was applied. Six motion parameters and average signal of grey matter, white matter, and cerebrospinal fluid were regressed out. Motion scrubbing was used to identify problematic volumes-bad volumes were labeled as a 1 and then used as an additional regressor. This technique was used so as to not remove continuous time which could lead to inaccurate estimates of the network state dynamics. Next, we extracted atlas-based preprocessed time series in standard space for each subject by calculating the mean of the preprocessed time series from each of the 36 Schaefer atlas regions (Schaefer et al. 2018) that make up the SN and DMN. These extracted time series were used as input for the MIND-Map toolbox (Khodaei et al. 2024) that uses a hidden semi-Markov model (HSMM) (Shappell et al. 2019) to estimate brain states and dynamics. Lastly, three timepoints were removed from the beginning and end of the signal to correct for edge effects caused by band-pass filtering.

### vi. Hidden Semi-Markov Modeling (HSMM)

We applied a HSMM to the 36 Schaefer atlas regions that make up the SN and DMN from our final sample of study participants. The list of regions used is in the Supplement Materials. While a detailed description of the HSMM is available in our previous work (Shappell et al. 2019), we provide a brief summary here. For each participant, we denote the ROI time series for each participant by *Y_i_*_1_*, …, Y_iT_*, where each *36*−dimensional vector *Y_it_* contains the BOLD measurements of the 36 ROIs at the *t^th^*timepoint for the *i^th^* participant. We assume that each observed vector, *Y_it,_* is generated by an unobserved (hidden) brain network state. This hidden state at time *t* for participant *i* is denoted by *S_it_*, where *S_it_* takes on discrete values. That is, *S_it_*∈ 1*…K*. Conditional on the current hidden state *S_it_*, the observed data *Y_it_* are modeled as following a multivariate normal distribution *Y_it_* ∼ N (*µ_s_,* Σ*_s_*), where the mean and covariance are dependent on the current (unknown) network states, for any *S* ∈ 1*, …, K*.

Thus, each network state is characterized by a unique pattern of mean activity across ROIs, along with a distinct covariance (and corresponding correlation) structure. We convert each state’s covariance matrix to a correlation matrix, which serves as the weighted brain network representation for that brain state. Moreover, the HSMM includes parameters that model (a) the transition probabilities between network states and (b) the dwell time distributions for how long each state persists after it is entered. These dynamic properties were then compared between the active and passive groups via permutation testing and mixed effects models (refer to Section v.).

For our analyses, we estimated a single set of network states using data pooled from all participants. The number of states must be defined in advance and is strongly influenced by such factors as the number of participants, timepoints, and regions of interest (ROIs). This is because increasing the number of states also increases the number of parameters that need to be estimated. To identify the optimal number of states, we ran the model using a range of state numbers (3-7) and evaluated the Euclidean distances between the resulting states. For each run, we calculated the minimum distance between any pair of states, with higher values indicating greater distinctiveness. The six-state model exhibited the highest minimum distance, suggesting it provided the most clearly differentiated set of states (see Supplementary Material for distance plots across runs). *add Moreover, for full details of the HSMM, including the complete data log-likelihood and parameter estimation routine, we direct readers to the Supplementary Material.

### vii. Statistical Inference and Permutation Testing

As described above, *a priori* analyses determined there were no significant differences between stress scores in either drinking state (normal versus abstained) immediately following the recovery period, so we focused our analyses on comparing state network dynamics between the active and passive recovery groups. Thus, each participant’s normal and abstained state recovery scan were included in the model to increase the number of parameters and allow us to include more ROIs for the analyses.

Each participant’s most probable sequence of true network states was estimated using the Viterbi Algorithm (Forney, 1973). The Viterbi algorithm takes the state estimates, transition probability estimates, dwell time density estimates, and initial state probability estimates obtained from fitting the HSMM on all participants and calculates each individual participant’s most likely sequence of states given those estimates and the participant’s BOLD time-series data. In other words, the Viterbi algorithm takes an individual’s time-series data and finds which of the six states estimated from the entire data set best fits that participant at each timepoint. Therefore, although the state estimates are based on the entire data set and are not subject-specific, when a state is active is subject-specific.

We examined the relationship between active/passive recovery groups and both occupancy times and empirical dwell time distributions of the brain states inferred from the HSMM. Occupancy time for each state was calculated by summing the total number of timepoints spent in that state, dividing by the length of the participant’s time-series, and then converting it to a percentage. Group averages for each state were calculated by finding the mean across all participants within each group. To calculate the empirical dwell time distribution for each state within a participant, we counted the number of consecutive timepoints spent in that state. These counts were then aggregated across all participants in each group, and a density estimate was computed for each group. For all dwell time calculations, we excluded the first and last state sequences for each participant to avoid introducing bias towards shorter sojourn times, as the exact duration of time spent in a state before or after the scan was unknown. After obtaining separate empirical sojourn distributions for each group across all six states, we compared the distributions between groups using Kullback-Leibler (KL) divergence, which measures the directed divergence between two probability distributions. A KL divergence of 0 indicates identical distributions, while a KL divergence of 1 suggests a substantial difference. For more details on KL divergence in this context, refer to Shappell et al. (2019). To assess whether any differences in occupancy time and/or dwell time distributions were statistically significant, we conducted a permutation test using 500 permuted samples.

Lastly, we investigated whether occupancy times or dwell time characteristics of the derived brain states were associated with stress scores. Specifically, we extracted occupancy time for each state, as well as the transition frequency into each state, for each participant. We fit mixed effects linear regression models, separately for each state occupancy time and transition frequency, with stress score as the outcome. Along with state occupancy time and transition frequency, an indicator variable for group membership (passive vs. active), an interaction term between group membership and the state occupancy time/transition frequency, as well as the average number of drinks consumed in the prior 30 days were included as predictors in the model. A random intercept was included to account for participants having 2 scans in the analysis. Stratified models were conducted for any models with a significant interaction. Therefore, a separate model was fit on the passive group and active group for any models that had a statistically significant interaction between group status and the state occupancy time/state transition frequency predictor.

### viii. Modularity Analysis for State Characterization

For each of the 6 brain states, we performed a modularity analysis to further understand each state’s defining topological characteristics. The modularity analysis identified community structures within each state. Modularity analysis estimates communities of nodes that have stronger within-group connections than their between-group connections (Newman, 2006). As a result, information flow is higher within communities compared to between communities. We applied the Newman spectral community detection method to the positive network of each state (Newman, 2013). A gamma value of 1 was used to divide the networks into more communities.

## III. Results

### i. Descriptive Statistics

Demographic and alcohol use characteristics for the sample can be found in Table 1. There was a total of 32 participants (19 female), 15 of which were assigned to the active recovery protocol while 17 were assigned to the passive recovery protocol. Of the full sample, 87.5% identified as white, 9.4% identified as African American or Black, 3.1% identified as Asian, and averaged 38.2 years of age. There were no significant differences between the active and passive group for the following demographic variables: age, race, sex, total years drinking, and percentage of drinking days in the past month. There was a significant difference in the average number of drinks on drinking days between the active and passive group (p=0.02), with the passive group averaging 2.5 drinks per day whereas the active group averaged 1.9 drinks per day. Therefore, we adjusted for this variable in our mixed effects models. In these models, the average number of drinks did not significantly predict stress scores. Additionally, there was a significant mean difference (p=0.03) in the 10-point stress scales taken immediately after the recovery period with participants in the mindfulness group reporting lower perceived stress (M=1.4, SD=1.36) than those is the passive, resting group (M=2.3, SD=1.52).

**Table 1.**
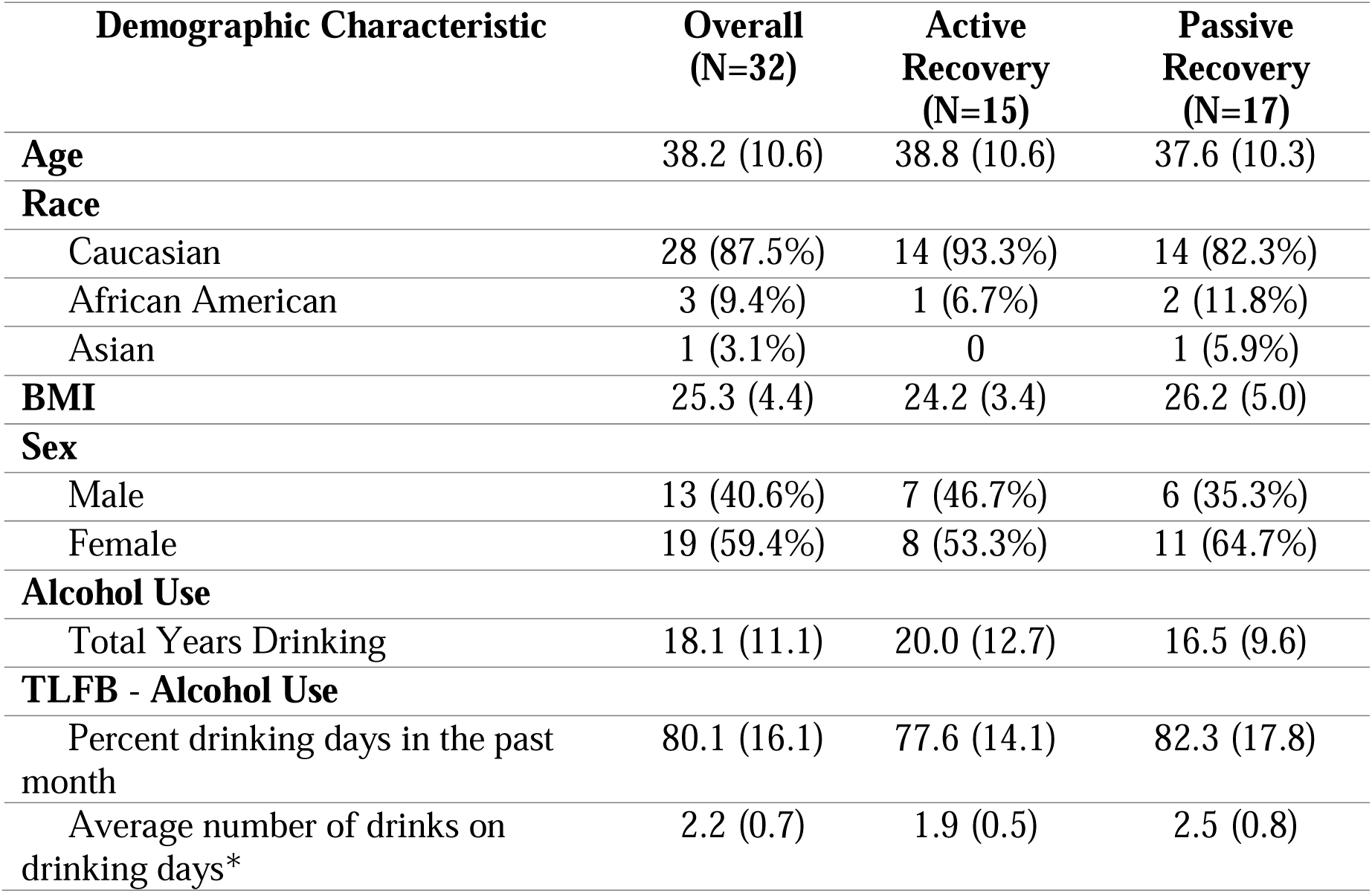
Sample characteristics listed as mean (SD) or frequency (percentage). *p<0.05 for a two-sample t-test comparing the average number of drinks on drinking days between the active group vs. the passive group.

### ii. Functional Brain State Characterization

A reference image of the DMN and SN as well as node community assignments, or modules, for each state can be found in Figures 2 and 3, respectively. The only state to have nodes belonging to the DMN and SN parcellated into two distinct modules was state 3. States 1, 2, and 5 had three modules while states 4 and 6 had four. For states 1 and 2, nodes of the SN remained in a module together, and the temporal regions of the DMN formed a module, shown in yellow. In state 5, the SN remained intact as well, but the DMN was split into two modules: one module consisting of bilateral temporal/parietal regions and the dorsolateral PFC in yellow and the other module contained the PCC and ventromedial PFC (vmPFC) in blue. In state 4, the SN remained intact, but the DMN formed three distinct modules: the prefrontal cortex (PFC) in blue, left and right temporal regions in green, and the PCC in yellow. In state 6, both the DMN and SN were split into four communities: anterior/posterior DMN in blue and yellow and anterior/posterior SN in red and green.

**Figure 2.**
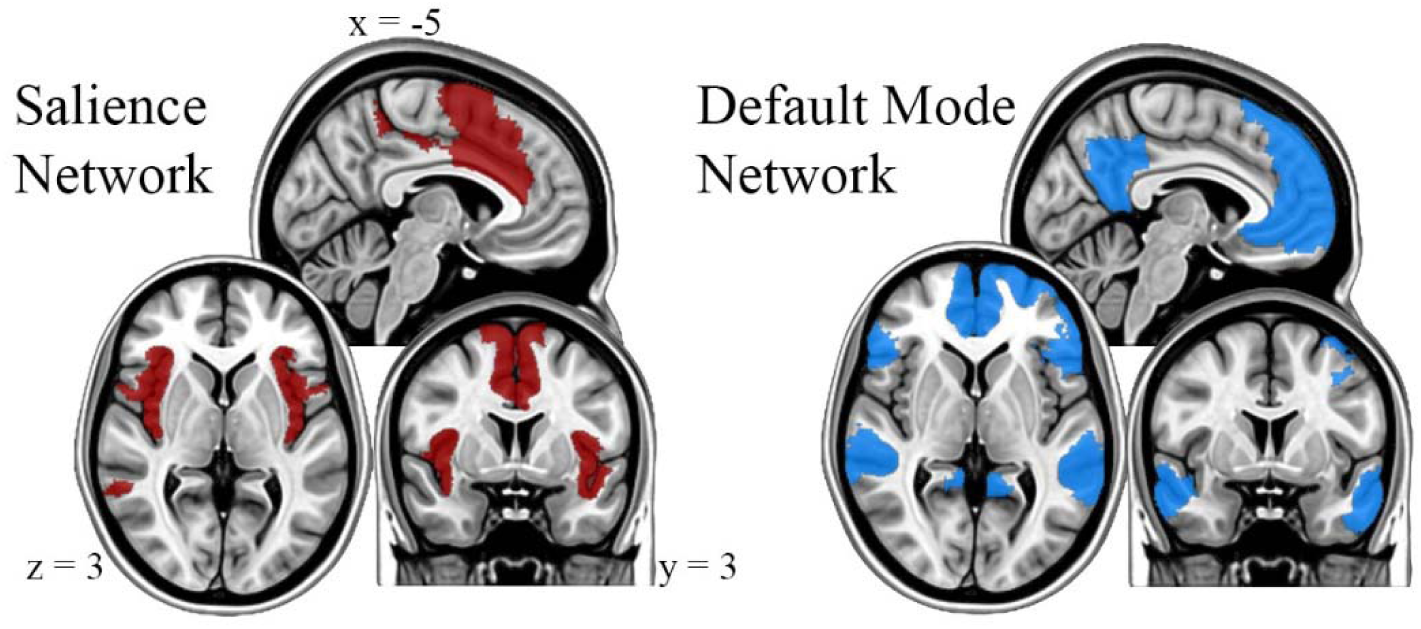
Localization of the salience network (SN, red) and default mode network (DMN, blue) as defined by the Schaefer 100-node atlas.

**Figure 3.**
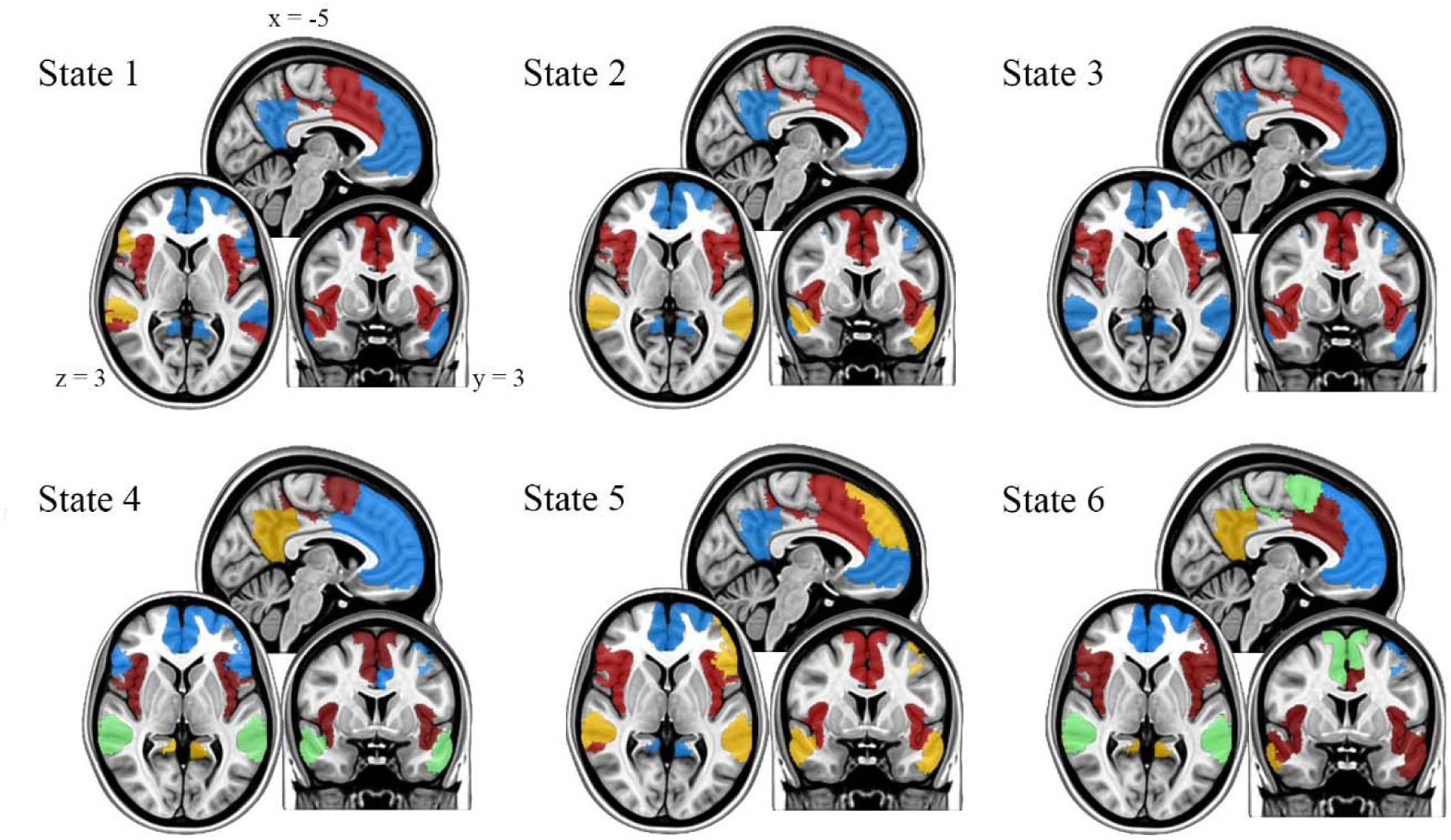
Module assignments for each state, represented by color grouping. Colors were matched as close as possible to the predefined SN and DMN. Differences in the color assignments in the modular maps across states reveal the “splitting” of nodes from either the DMN or SN.

Mean activity maps are represented by the mean z-scored BOLD signal for nodes in each state compared to activity in the other states (Figure 4). In state 1, anterior DMN nodes are primarily active, including the left prefrontal cortex and temporal/parietal regions. States 3, 5, and 6 are characterized by high posterior DMN activity. Specifically, the posterior cingulate cortex, a hub of the DMN, displays high activity across all three states. In state 2, many SN nodes show high activity, specifically the insula and the dorsal anterior cingulate cortex (dACC), both of which are hubs within the SN. Finally, state 4 is characterized mainly by high activity in the bilateral insula.

**Figure 4.**
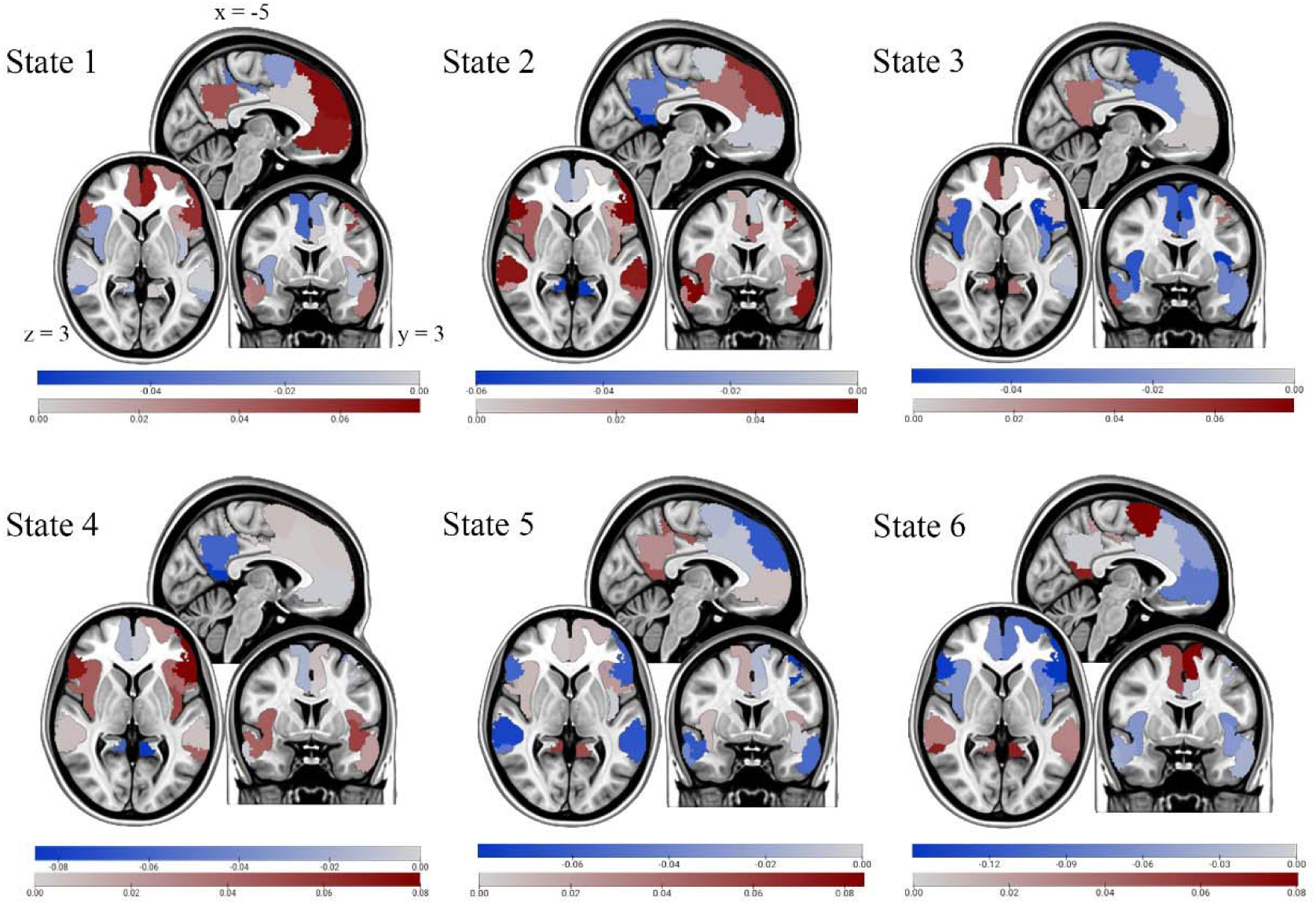
Mean activation maps for each state. Dark blue colors indicate lower activity, grey/white colors indicate average activity, and dark red colors indicate higher activity across states. Activity is relative to that observed during the other states, not relative to a baseline.

### iii. Occupancy Time, Transition Frequency, and Dwell Time

Participants in the active mindfulness group spent significantly more total scan time, on average, in state 2 during the recovery period (p=0.0002) compared to the passive recovery group. Furthermore, active recovery participants transitioned significantly more frequently into state 2 (p=0.0002) throughout the recovery period compared to participants in the passive recovery group. Conversely, participants in the passive group spent significantly more time in states 1 and 3 during the recovery period (p=0.0002, p=0.0002) and had significantly more transitions into states 1 and 3 (p=0.0002, p=0.0002) compared to the active recovery group. Results of the permutation test for each state and each state characteristic can be found in Table 2. Bar plots depicting occupancy time and transition frequency for each cohort in each state can be seen in Figure 5.

**Figure 5.**
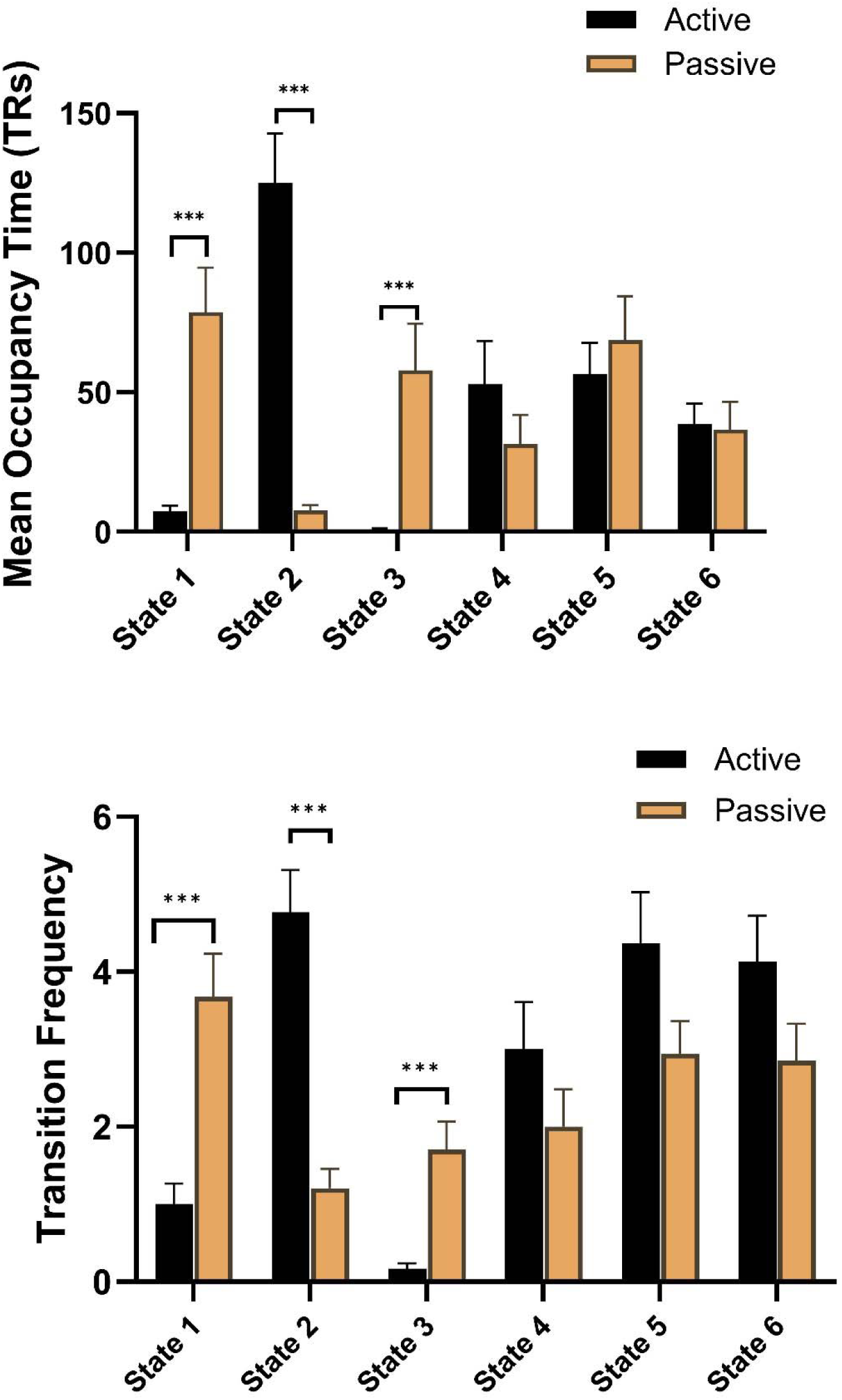
(Top) Mean occupancy time, represented by average number of TRs, for each cohort in each state. (Bottom) Transition frequency for each cohort to each state. ***p<0.001.

**Table 2.** Permutation test results for each state and each dynamic property. *p<0.05 for a permutation test comparing active vs. passive groups.

| State | Occupancy Time | Sojourn/Dwell Time | Transition Frequency |
| --- | --- | --- | --- |
| 1 | **0.0002\*** | **<0.0001\*** | **0.0002\*** |
| 2 | **0.0002\*** | **<0.0001\*** | **0.0002\*** |
| 3 | **0.0002\*** | **0.004\*** | **0.0002\*** |
| 4 | 0.2416 | 0.858 | 0.1129 |
| 5 | 0.5461 | 0.406 | 0.6335 |
| 6 | 0.8897 | 0.29 | 0.3766 |

Dwell (sojourn) time represents how many consecutive time points a participant spends in a state before transitioning to another. There was a significant difference in sojourn distributions between the two groups for states 1, 2, and 3 (p<0.0001, p<0.0001, p=0.004). Specifically, we found that individuals in the active recovery group had shorter dwell times in states 1 and 3 and a longer dwell time in state 2 before switching to another state. Participants in the passive recovery group had a shorter dwell time in state 2 and a longer dwell time in states 1 and 3 before switching to another state. Sojourn distributions for the six states for each recovery group can be found in Figure 6. Higher curves to the left of the x-axis represents shorter dwell times for that state.

**Figure 6.**
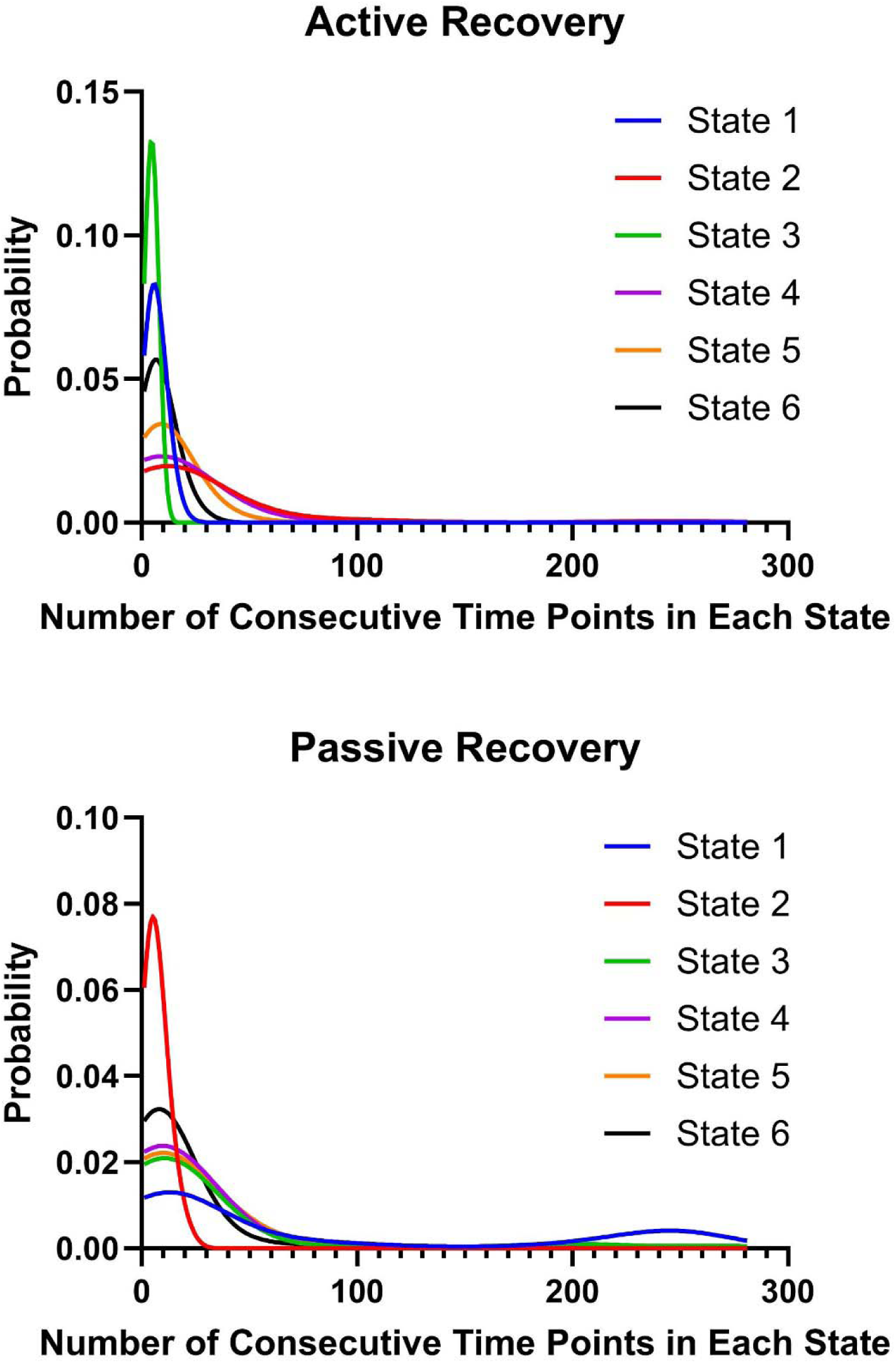
Estimated dwell time distributions for each state for the active (top) and passive (bottom) recovery groups.

### iv. Mixed Effects Model & Stress Scores

Results from the repeated measures mixed effects model determined there was an association between brain state dynamics and stress scores immediately following the recovery period. Interestingly, we found that increased occupancy time in state 2 was significantly associated with lower stress scores in the passive group (p=0.002) following the recovery period (Table 3). No other associations between state occupancy time and stress scores, as well as transition frequency to the other 5 states and stress scores, were found. Additionally, there were no significant effects found when examining the association between total transition frequency (across all 5 states) and stress scores.

**Table 3.**
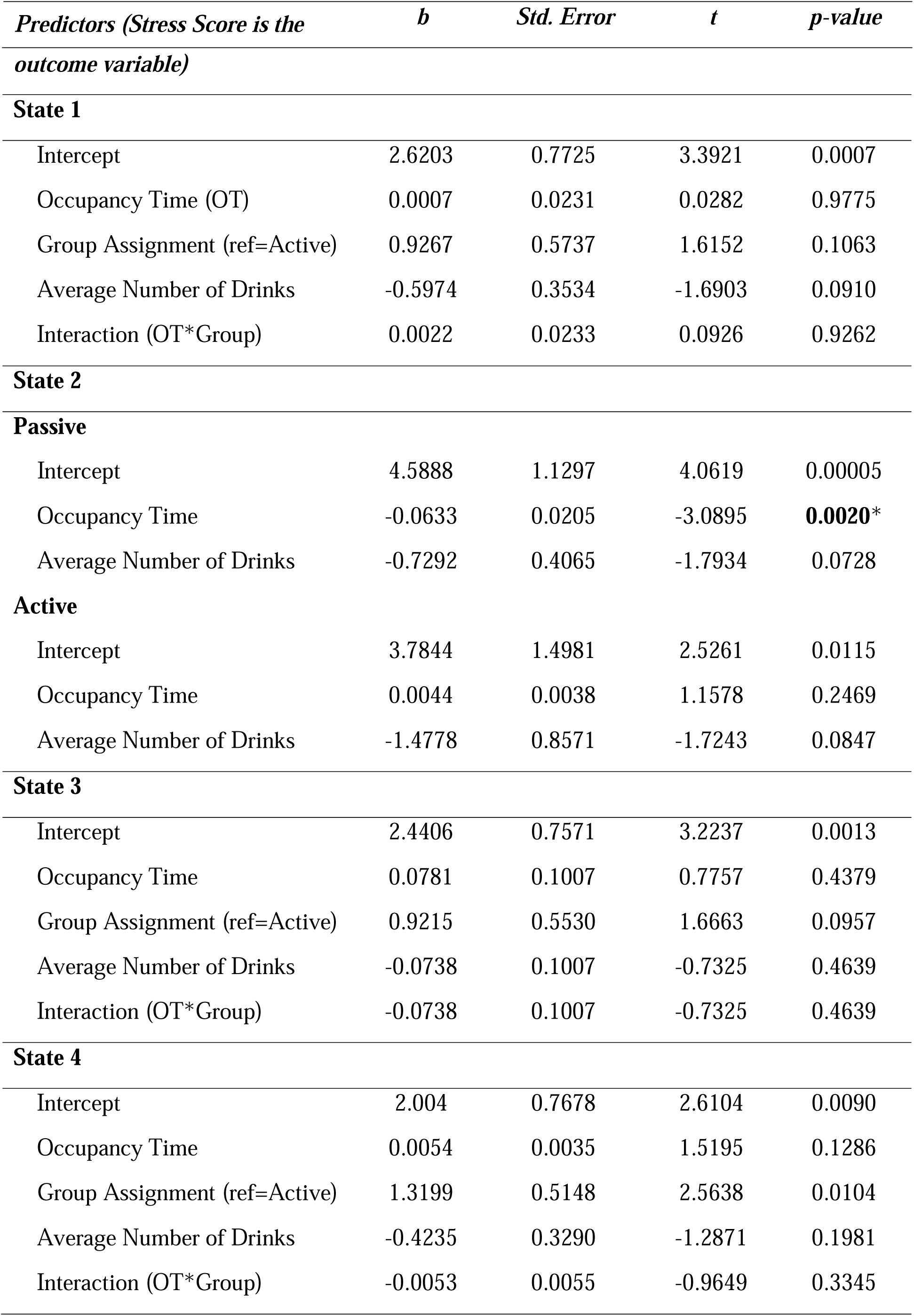
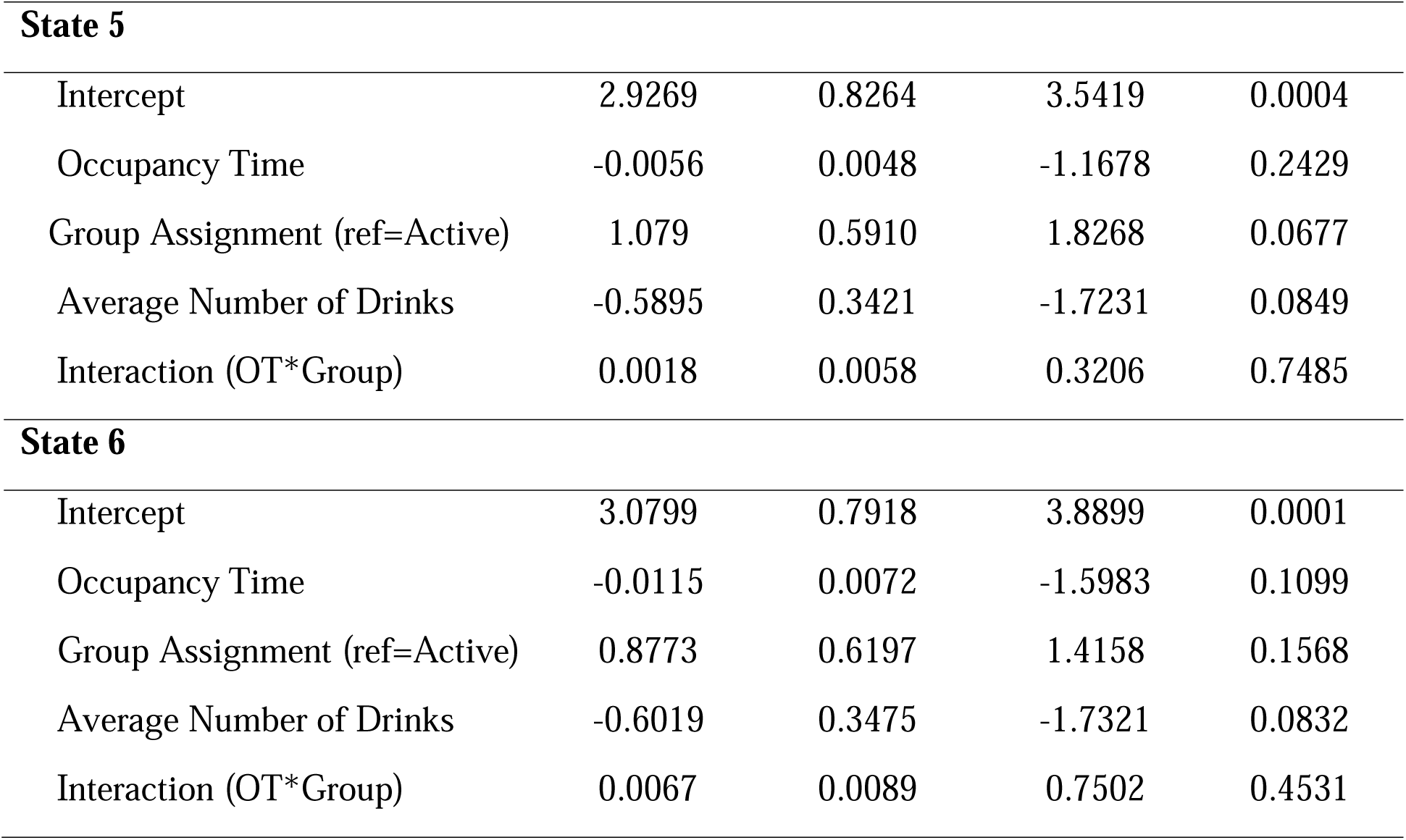
Results of mixed effect model. State 2 had a significant interaction for state occupancy time, so a stratified analysis was performed on the active and passive groups separately. *p<0.05

## IV. Discussion

Following exposure to an individualized acute stressor, we contrasted the effects of a brief, guided mindful-based recovery period to resting control on the dynamic functional connectivity of the DMN and SN in individuals who self-reported moderate to heavy alcohol consumption. We determined optimal state(s) for both the guided mindfulness (active recovery) group and the control (passive recovery) group and investigated how each group moved through these states. Additionally, we determined which dynamic characteristics were associated with either higher or lower stress scores for each group. Specifically, we found that 6 optimal states emerged from a dynamic analysis of brain states during the recovery period supporting our hypotheses that, compared to passive recovery, guided mindfulness yielded higher occupancy time and transition frequency groups to states in which nodes of the SN were more active; conversely, the control group exhibited higher occupancy time and transition frequency to states in which nodes of the DMN were more active. We also observed lower perceived stress following the recovery period among participants in the mindfulness group as compared to the control group, and a significant correlation between occupancy time and stress scores following the recovery period in the passive group. Below, we discuss possible interpretations of our findings.

The active recovery group had spent longer periods of time, consecutively and on average, in state 2. Moreover, individuals in the active group had transitioned into this state more often than the passive group. Modularity analysis of state 2 revealed unique patterns of functional connectivity, with three distinct modules emerging. Nodes of the SN formed a module together, temporal nodes of the DMN formed another, and the PFC and PCC formed a third module. Activity maps revealed increased activity in the SN module, the temporal DMN module, and the dorsomedial PFC (dmPFC), part of the larger PFC-PCC module, whereas the remainder of the PFC-PCC module showed low-to-normal activity relative to the other 5 states. The fact that participants in the active group spent more time in state 2 was not surprising, as several studies have implicated increased activity of the SN as a key mechanism underlying positive therapeutic effects of mindfulness-based interventions (Bremer et al 2022; Doll et al. 2015; Hasenkamp and Barsalou, 2012). The fact that participants in the passive group spent more time in states characterized by increased activity in modules of the DMN implicated in rumination offers strong support for the beneficial role of mindfulness-based recovery.

Interestingly, as noted in the modularity analysis, there were also two distinct modules of the DMN that were active in state 2: the temporal nodes of the DMN and the dmPFC. Recent work has observed increased activity in cortical regions of the temporal lobe of the DMN, specifically within the context of processing information related to the self (Ribeiro Da Costa et al. 2022). While this may seem counterintuitive to the positive effects of mindfulness, it is important to emphasize that paying attention to and processing interoceptive cues, which was a central feature of the active coping script, is known to enhance a more integrated perspective of the self via the right hemisphere (Johnstone et al. 2021), and is the basis for mindful awareness body therapy that was designed specifically to address dysfunction in emotional regulation among participants with alcohol abuse disorders (Price & Hooven, 2018). Additionally, the increase in dmPFC activity seen in state 2 is consistent with findings showing an increase in dmPFC activity during meditation, most likely because of its role in cognitive emotional evaluation and regulation (Brewer et al. 2011; Doll et al. 2015).

Hence, we conclude that the active recovery condition—specifically increased activity in nodes of the temporal DMN, dmPFC, and SN—were working in concert to maintain an integrated sense of self in present moment awareness (Brewer et al. 2011). It is known that allocating attention to interoceptive cues is dependent upon the SN (Bremer et al. 2022; Doll et al. 2015; Hasenkamp and Barsalou, 2012), plays a major role in the right hemisphere’s construction of an integrated sense of self (Johnstone et al. 2021), is a central feature of mindful-based training programs (A. Niazi & S. Niazi, 2011), and likely plays a role in the experience of mindful-based transcendent states (Handley et al. 2020).

States 1 and 3 were significant for the passive recovery group, in that subjects transitioned more often into these states and spent longer consecutive and overall timepoints in these two states compared to the active recovery group. Modularity analysis for state 3 determined that nodes of the SN formed a module together while nodes of the DMN assembled into another module. In this state, the PCC, a hub of the DMN, exhibited increased activation relative to the other states. State 1 connectivity patterns were unique in that most nodes of the PFC, PCC, and left temporal nodes of the DMN were in a module together, nodes of the SN were in another module, and the right temporal and ventral PFC (vPFC) were in a third module. Activation maps for state 1 were characterized by increased activity of the PCC and PFC while state 3 activity was localized to the PCC only, relative to the remaining states. Increased activity of hubs within the DMN and decreased activity of the SN, as seen in states 1 and 3, is consistent with previous work establishing the role of this subnetwork as being active during periods of rest and may even be implicated in mind-wandering and rumination (Mason et al. 2007). Further, this supports previous work that has observed these two subnetworks exhibiting anticorrelated activity at rest (Anticevic et al. 2012). As described above, the passive recovery period was established as a rest period to be used as a comparison to active recovery. Thus, as hypothesized, the passive group spent more time engaged in DMN activity known to be associated with rumination and less time in SN activity relative to the mindful group.

## V. Limitations

The current study has some limitations to note. First, the mindfulness session used in this study was a one-time, 10-minute, guided experience with no training specific to the intervention prior to the study. Although single bouts of mindfulness have proved successful in reducing stress and stress-related alcohol consumption and craving (Kamboj et al. 2017; Shuai et al. 2020), studies have shown that the more experienced one is in meditation, the more likely they are to build resilience to stress and be more successful in coping with stress in the future (Zgierska et al. 2008). Hence, an important extension of this study would be to expose participants to formal training in mindfulness, to have them employ skills in response to acute stressors across the day, and to evaluate the effects of this training using both experience sampling methodology and controlled study in a laboratory setting.

Second, our analysis estimated that six states were the best fit for the HSMM model, as these six states produced the most stable state sequences across the sample. However, participants may be entering more (or fewer) than six states during a 10-minute mindfulness session scan. Further, ROIs used were cortical nodes of the SN and DMN, thus possibly missing important contributions of subcortical structures and the central executive network (CEN), which has also been a network of interest during mindfulness (Doll et al. 2015). Future methodological work in this area should allow for larger state sizes (i.e., more brain regions to be included in the analyses), as well as a way to quantify whether the number of and estimate of the network states fits each individual well.

Lastly, we employed a single item of perceived stress following the active and passive recovery conditions. In hindsight, it would have been beneficial to have more comprehensive assessments of stress and affect following recovery. We highly recommend these additional assessments in future study designs.

## VI. Conclusion

To our knowledge, this is the first study to evaluate how mindfulness-based techniques following an acute stressor in moderate to heavy drinkers influence brain networks employing dynamic connectivity methodology—HSMM. The use of HSMM enabled us to track changes in brain states between our experimental conditions that would have been masked by commonly used “static” methodology. Specifically, guided mindfulness yielded higher occupancy time and transition frequency groups to states in which nodes of the SN were more active, whereas the control group exhibited higher occupancy time and transition frequency to states in which nodes of the DMN associated with rumination were more active. We also observed lower perceived stress following the recovery period among participants in the mindfulness group as compared to the control group.

The results of the current study on mindful coping following an acute stressor are the first to demonstrate this pattern of network connectivity, albeit in the context of a single, brief mindful-based intervention. A question of both conceptual and practical relevance is whether mindfulness-based training enables participants to achieve similar brain states when confronted with life stress as well as the features of training that maximize treatment efficacy.

## Data and Code Availability

The data presented in this study is not publicly available. The toolbox and associated code can be made available upon request.

## Informed Consent

Informed consent was obtained from all participants involved in this study.

## Institutional Review Board Statement

This study was conducted in accordance with the guidelines of the Declaration of Helinski and approved by the Institutional Review Board of Wake Forest Health Sciences (IRB00028739) on 7/15/2014.

## Acknowledgements

We thank the participants for their time and participation as well as Dr. Rhiannon Mayhugh for all of her work on this project.

