## Supplemental Information for "The Effects of Mindfulness on Brain Network Dynamics Following an Acute Stressor in a Population of Moderate to Heavy Drinkers"

**Schaefer Atlas ROIs and Subnetwork Assignment**

| **ROI Label** | **Subnetwork Assignment** |
| --- | --- |
| 38 | DMN |
| 39 | DMN |
| 40 | DMN |
| 41 | DMN |
| 42 | DMN |
| 43 | DMN |
| 44 | DMN |
| 45 | DMN |
| 46 | DMN |
| 47 | DMN |
| 48 | DMN |
| 49 | DMN |
| 50 | DMN |
| 90 | DMN |
| 91 | DMN |
| 92 | DMN |
| 93 | DMN |
| 94 | DMN |
| 95 | DMN |
| 96 | DMN |
| 97 | DMN |
| 98 | DMN |
| 99 | DMN |
| 100 | DMN |
| 24 | SN |
| 25 | SN |
| 26 | SN |
| 27 | SN |
| 28 | SN |
| 29 | SN |
| 30 | SN |
| 74 | SN |
| 75 | SN |
| 76 | SN |
| 77 | SN |
| 78 | SN |

Table 1. ROIs used for the current study as well as the subnetwork assignment. The atlas used was the Schaefer 100-node 7 network atlas. DMN = Default Mode Network, SN = Salience Network.

**Further Information on the Hidden Semi-Markov Model (HSMM)**

Our previously validated HSMM framework for inferring time-varying brain networks from fMRI data estimates a set of network states, dwell/sojourn time distributions, and probabilities for switching from one state to another – all at the group level (Shappell et al. 2019). Individual estimates of state sequences can be further derived by running a Viterbi algorithm (Forney, 2005), and those state sequences can be used to obtain individual estimates of overall time spent in each state, as well as transition probabilities. Briefly, we denote the ROI time-series data for each participant by $Y_{i1}, \ldots, Y_{iT}$, where each $p$−dimensional vector $Y_{iT}$ contains the BOLD measurements of the $p$ROIs at the $t^{th}$ timepoint for the $i^{th}$ participant. The collection of vectors of the observed time-series data is denoted by $\tilde{Y}_{i}$. Now suppose a unique brain network state gives rise to each $Y_{it},$ but we cannot observe this. We represent this hidden network index variable underlying the observed time-series vectors at a particular observation time by $S_{it},$where $S_{it}$ takes on discrete values. That is, $S_{it}, \in\left\{ 1,\ldots, K \right\}.$The vector of hidden network state variables, $S_{i1}, \ldots, S_{iT}$, for a single individual is denoted by $\tilde{S}_{i}$ . We assume that each $Y_{it}$follows a multivariate Gaussian distribution $Y_{it} \sim N\left( \mu_{s}, \Sigma_{s} \right),$ where the mean and covariance depend on the current (unknown/hidden) network state. Thus, each network state has unique mean activations across ICs, and its own covariance structure between ICs. The complete data log-likelihood of the HSMM for one participant, assuming $K$ unique network states, can be written as:

$$\log P(\tilde{Y}_{i}=\tilde{y}_{i},\tilde{S}_{i}=\tilde{s}_{i};\mu_{1:K,}\Sigma_{1:K}, P,d_{1:K})=log f\left( \tilde{s}_{i},\tilde{y}_{i} \right)$$

$=\log f\left( \tilde{y}_{i}|\tilde{s}_{i} \right)+\log f\left( \tilde{s}_{i} \right)$

$=\sum_{t=1}^{T} \log f\left( y_{it} | s_{it} \right)+ \sum_{r=2}^{R} \log(f(s_{ir}|s_{i(r-1)}) d_{s_{ir}}(u_{ir}))+$

$$\log(f(s_{iR}|s_{i(R-1)}) D_{s_{iR}}(u_{iR})) \log(f\left( s_{i1} \right)d_{s_{i1}}(u_{i1})),$$

where $R$ = total number of network state changes + 1, $f\left( s_{1} \right)$ is the probability of starting in a particular state, $d(u)$ is the sojourn time density, $s_{r}$ is the $r^{th}$ visited state, and $u_{r}$is the time spent in that state. The first term is based on the conditional distribution of the observed BOLD signal vector given the underlying hidden network, which takes on a Gaussian distribution, as noted above. The second portion of the equation is made up of two parts. The first is a transition probability matrix, denoted $P$, where the probability at row q and column v represents the probability of transitioning from network state $q$ to $v$ (i.e., $p_{qv}=P(s_{r}=v, s_{r-1}=q$)). The second part, $d_{s_{r}}\left( u_{r} \right),$ represents the dwell time/sojourn distribution. The third portion accounts for the last state a participant enters. Note that

$D\left( u \right)= \sum_{V\geq u} d(v)$

is the survivor function and pertains only to the sojourn time in the final state. It allows us to avoid the assumption that the process is leaving the final state immediately after time $T.$ Lastly, the fourth term accounts for an individual’s initial network state. The log-likelihood is then summed across subjects, and a Maximization Expectation (E-M) algorithm is used to estimate the model parameters.

**Minimum State Distance Plot**

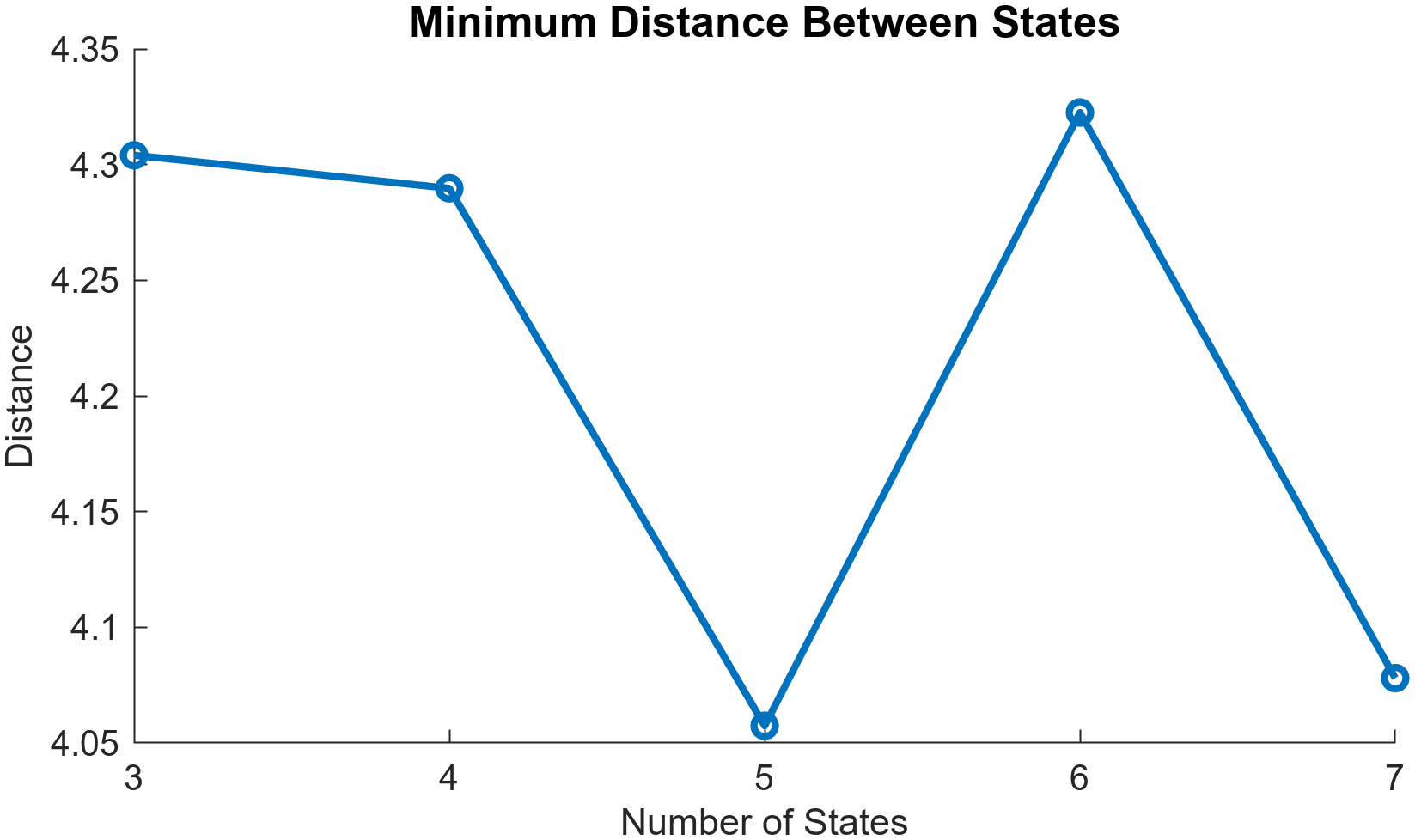

Figure 1. Plot depicting the minimum distance between state pairs for each run. The run with the highest minimum distance between state pairs specifies the optimal number of states. In this study, our optimal number of states was 6.

**HSMM Results with Outlier Included**

One participant was removed from the study due to having a high number of transitions at the beginning of each scan, likely due to motion, and causing a large influence on between-group differences for occupancy time, transition frequency, and dwell time. Additionally, including this participant resulted in no significant associations between stress scores and brain state dynamics. Results of the HSMM with the outlier included are described below.

Permutation testing revealed that participants in the active group spent more time, on average, in states 2 and 4 compared to the passive group (p=0.0002, p=0.0086). In contrast, the passive group spent more time, on average, in states 1 and 3 compared to the active group (p=0.0002, p= 0.0024). Additionally, the active group had significantly higher transitions into states 2 and 4 (p=0.0002, p=0.0092), whereas the passive group had more transitions into states 1 and 3, comparatively (p=0.0002, p=0.0002). Finally, the active group had a significantly longer dwell time in state 2 (p=0) while the passive group had a significantly longer dwell time in states 1 and 6 (p=0.002, 0.022). Lastly, there were no significant associations detected between state dynamics and stress scores following recovery ( all p’s>0.05).

Notably, the state module and state activation maps did not differ greatly. In the original dataset, states 1, 2, and 5 had three modules, states 4 and 6 had four, and state 3 had two. When including the outlier in the dataset, states 1, 2, 4, and 5 had three modules, state 3 had two, and state 6 had four. Activity maps for states 1, 4, and 5 are characterized by high anterior and posterior DMN activity while state 2 is characterized by high SN activity. State 3 is characterized by high posterior DMN activity, mainly the PCC. Finally, state 6 is characterized by high activation of SN nodes along the midline. For a full list of results of the permutation tests, mixed effects model, subject state sequences, state module and activity maps, all with the outlier included, see below.

| State | Occupancy Time | Sojourn/Dwell Time | Transition Frequency |
| --- | --- | --- | --- |
| 1 | **0.0002*** | **0.008*** | **0.0002*** |
| 2 | **0.0002*** | **<0.0001*** | **0.0002*** |
| 3 | **0.0002*** | 0.076 | **0.0002*** |
| 4 | **0.0006*** | 0.11 | **0.0072*** |
| 5 | 0.3657 | 0.438 | 0.4989 |
| 6 | 0.6663 | **0.014*** | 0.7735 |

Table 2. Permutation test results for each state and each dynamic characteristic with the original outlier included. *p<0.05 for a permutation test comparing active vs. passive groups

| ***Predictors (Stress Score is the outcome variable)*** | ***b*** | ***Std. Error*** | ***t*** | ***p-value*** |
| --- | --- | --- | --- | --- |
| **State 1** | | | | |
| Intercept | 2.4094 | 0.7993 | 3.0142 | 0.0026 |
| Occupancy Time (OT) | -0.0142 | 0.0194 | -0.7305 | 0.4651 |
| Group Assignment (ref=Active) | 0.7482 | 0.5690 | 1.3149 | 0.1886 |
| Average Number of Drinks | -0.4242 | 0.3535 | -1.1999 | 0.2302 |
| Interaction (OT*Group) | 0.0161 | 0.0197 | 0.8174 | 0.4137 |
| **State 2** | | | | |
| Intercept | 1.9363 | 0.8138 | 2.3793 | 0.0173 |
| Occupancy Time (OT) | 0.0029 | 0.0031 | 0.9161 | 0.3596 |
| Group Assignment (ref=Active) | 1.5082 | 0.6398 | 2.3573 | 0.0184 |
| Average Number of Drinks | -0.4068 | 0.3454 | -1.1779 | 0.2389 |
| Interaction (OT*Group) | -0.0263 | 0.0215 | -1.2249 | 0.2206 |
| **State 3** | | | | |
| Intercept | 1.9553 | 0.7858 | 2.4885 | 0.0128 |
| Occupancy Time (OT) | -0.0068 | 0.0535 | -0.1265 | 0.8993 |
| Group Assignment (ref=Active) | 0.6269 | 0.5659 | 1.1077 | 0.2679 |
| Average Number of Drinks | -0.2453 | 0.3595 | -0.6823 | 0.4950 |
| Interaction (OT*Group) | 0.0115 | 0.0536 | 0.2154 | 0.8295 |
| **State 4** | | | | |
| Intercept | 1.9842 | 0.7900 | 2.5116 | 0.0120 |
| Occupancy Time (OT) | 0.0021 | 0.0039 | 0.5276 | 0.5978 |
| Group Assignment (ref=Active) | 1.2039 | 0.5922 | 2.0329 | 0.0421 |
| Average Number of Drinks | -0.3447 | 0.3531 | -0.9763 | 0.3289 |
| Interaction (OT*Group) | -0.0055 | 0.0068 | -0.8091 | 0.4185 |
| **State 5** | | | | |
| Intercept | 2.3906 | 0.8037 | 2.9745 | 0.0029 |
| Occupancy Time (OT) | -3.2228 | 0.0041 | -0.7877 | 0.4309 |
| Group Assignment (ref=Active) | 1.0660 | 0.5779 | 1.8443 | 0.0651 |
| Average Number of Drinks | -3.8875 | 0.3477 | -1.1182 | 0.2635 |
| Interaction (OT*Group) | -4.8070 | 0.0050 | -0.0001 | 0.9999 |
| **State 6** | | | | |
| Intercept | 2.6035 | 0.7889 | 3.2999 | 0.0009 |
| Occupancy Time (OT) | -0.0103 | 0.0074 | -1.3949 | 0.1630 |
| Group Assignment (ref=Active) | 0.7545 | 0.5727 | 1.3174 | 0.1877 |
| Average Number of Drinks | -0.4049 | 0.3375 | -1.1998 | 0.2302 |
| Interaction (OT*Group) | 0.0078 | 0.0085 | 0.9127 | 0.3614 |

Table 3. Results of mixed effect model in which occupancy time is a predictor of stress scores (with original outlier included). No significant interactions were found, so there was no stratified analysis performed.

| ***Predictors (Stress Score is the outcome variable)*** | ***b*** | ***Std. Error*** | ***t*** | ***p-value*** |
| --- | --- | --- | --- | --- |
| **State 1** | | | | |
| Intercept | 2.4287 | 0.7863 | 3.0888 | 0.0020 |
| Transition Frequency (TF) | -0.1347 | 0.1388 | -0.9707 | 0.3317 |
| Group Assignment (ref=Active) | 0.8456 | 0.5839 | 1.4480 | 0.1476 |
| Average Number of Drinks | -0.4058 | 0.3449 | -1.1764 | 0.2394 |
| Interaction (TF*Group) | 0.1307 | 0.1606 | 0.8139 | 0.4157 |
| **State 2** | | | | |
| Intercept | 2.7969 | 0.9091 | 3.0767 | 0.0021 |
| Transition Frequency (TF) | -0.0946 | 0.0845 | -1.1205 | 0.2625 |
| Group Assignment (ref=Active) | 0.6447 | 0.6738 | 0.9569 | 0.3386 |
| Average Number of Drinks | -0.4387 | 0.3482 | -1.2601 | 0.2076 |
| Interaction (TF*Group) | 0.0099 | 0.1885 | 0.0527 | 0.9579 |
| **State 3** | | | | |
| Intercept | 2.0342 | 0.7706 | 2.6396 | 0.0083 |
| Transition Frequency (TF) | -0.1162 | 0.3200 | -0.3629 | 0.7166 |
| Group Assignment (ref=Active) | 0.5592 | 0.5601 | 0.9984 | 0.3181 |
| Average Number of Drinks | -0.2707 | 0.3512 | -0.7708 | 0.4408 |
| Interaction (TF*Group) | 0.2973 | 0.3356 | 0.8859 | 0.3757 |
| **State 4** | | | | |
| Intercept | 2.6059 | 0.7907 | 3.2958 | 0.0009 |
| Transition Frequency (TF) | -0.0779 | 0.0793 | -0.9827 | 0.3258 |
| Group Assignment (ref=Active) | 1.1089 | 0.5779 | 1.9191 | 0.0549 |
| Average Number of Drinks | -0.4442 | 0.3324 | -1.3368 | 0.1814 |
| Interaction (TF*Group) | -0.1593 | 0.1335 | -1.1934 | 0.2327 |
| **State 5** | | | | |
| Intercept | 2.5893 | 0.7780 | 3.3281 | 0.0009 |
| Transition Frequency (TF) | -0.0841 | 0.0721 | -1.1667 | 0.2433 |
| Group Assignment (ref=Active) | 1.0169 | 0.5914 | 1.7198 | 0.0855 |
| Average Number of Drinks | -0.4344 | 0.3295 | -1.3182 | 0.1874 |
| Interaction (TF*Group) | -0.0129 | 0.1195 | -0.1078 | 0.9142 |
| **State 6** | | | | |
| Intercept | 2.8775 | 0.8082 | 3.5602 | 0.0004 |
| Transition Frequency (TF) | -0.1493 | 0.0849 | -1.7596 | 0.0785 |
| Group Assignment (ref=Active) | 0.7255 | 0.6272 | 1.1568 | 0.2473 |
| Average Number of Drinks | -0.4452 | 0.3345 | -1.3307 | 0.1833 |
| Interaction (TF*Group) | 0.0710 | 0.1174 | 0.6047 | 0.5454 |

Table 4. Results of mixed effect model in which transition frequency is a predictor of stress scores (with original outlier included). No significant interactions were found, so there was no stratified analysis performed.

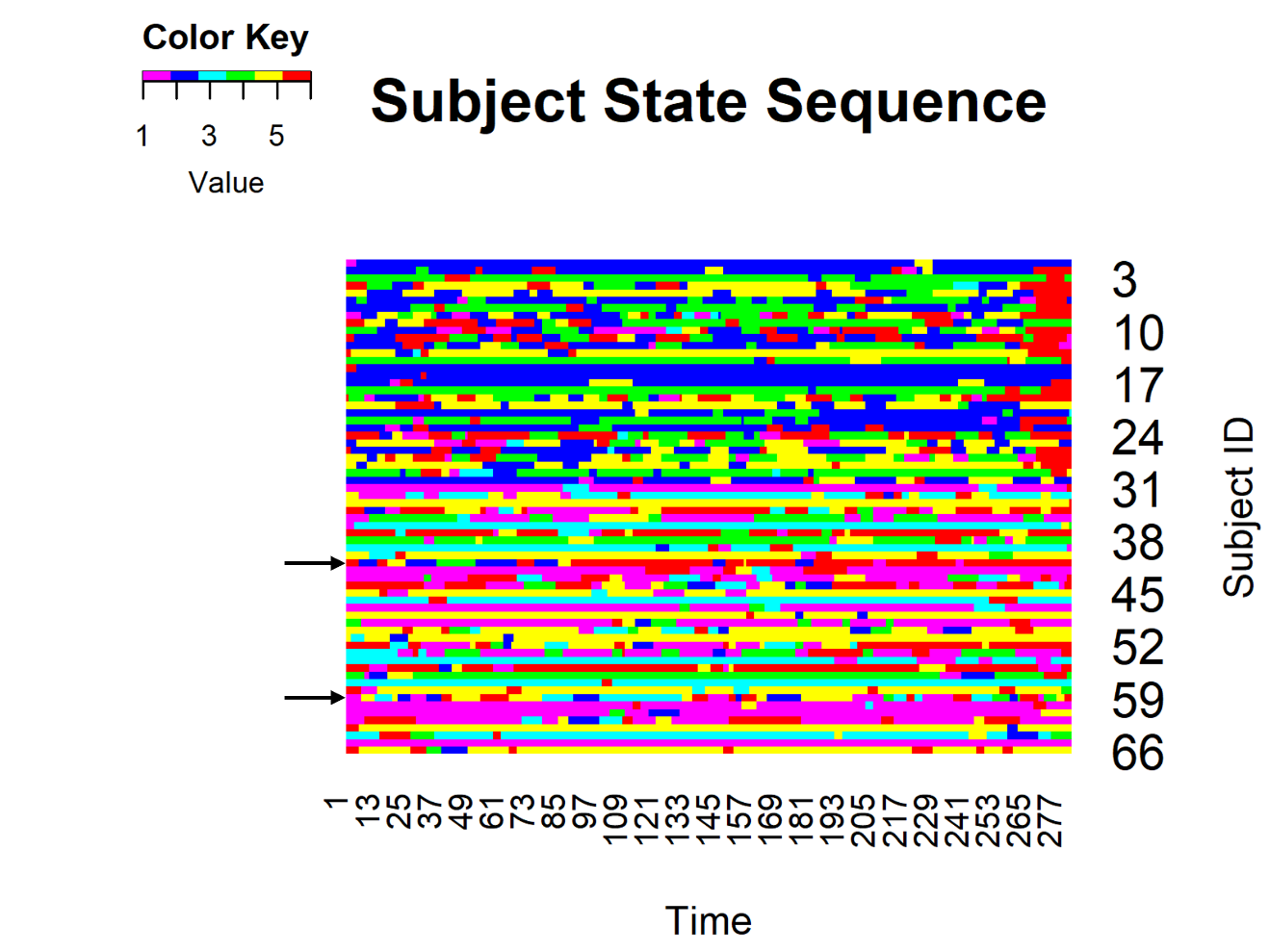
Figure 2. Subject state sequences of both groups during their active/passive recovery scan. The state sequences of the outlier participant are labeled with black arrows. Both scans have a high number of transitions at the beginning of the scan.

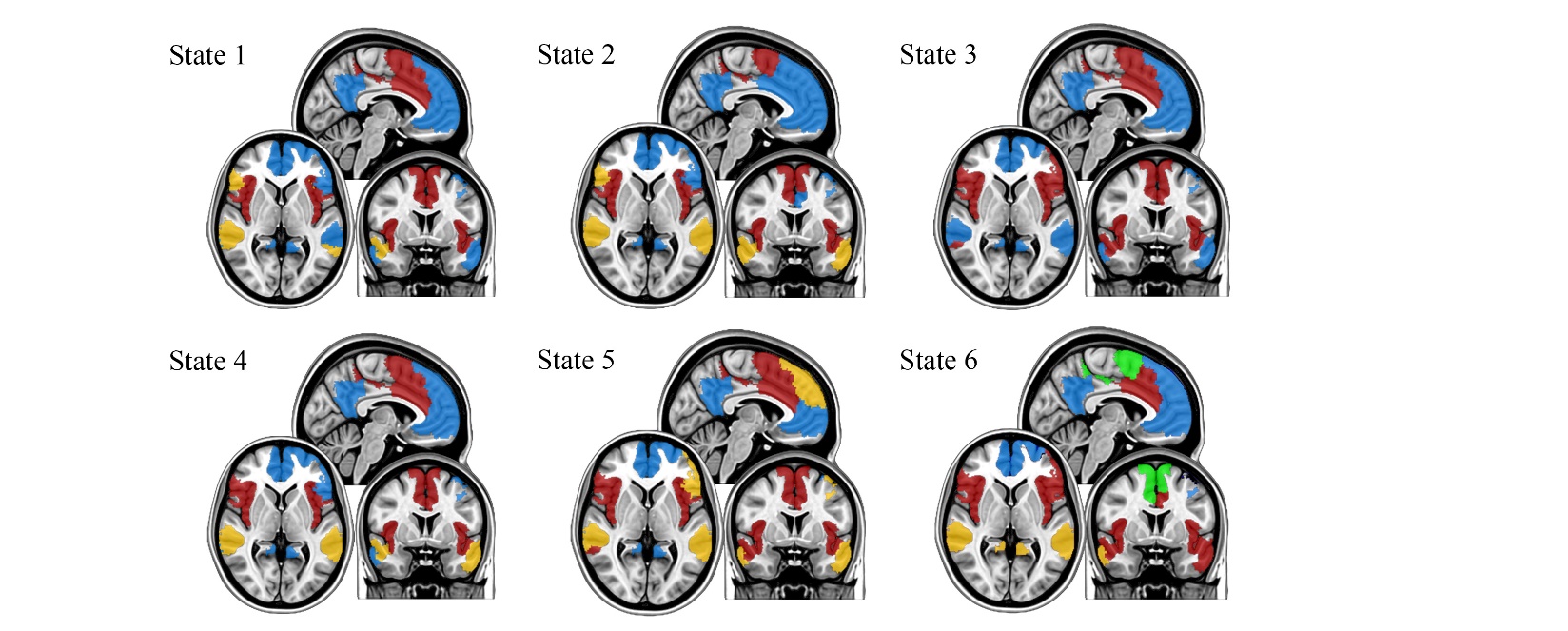
Figure 3. Module maps for each of the six states with the outlier included, represented by color grouping. Colors were preserved across states, to easily observe module assignments across each state.

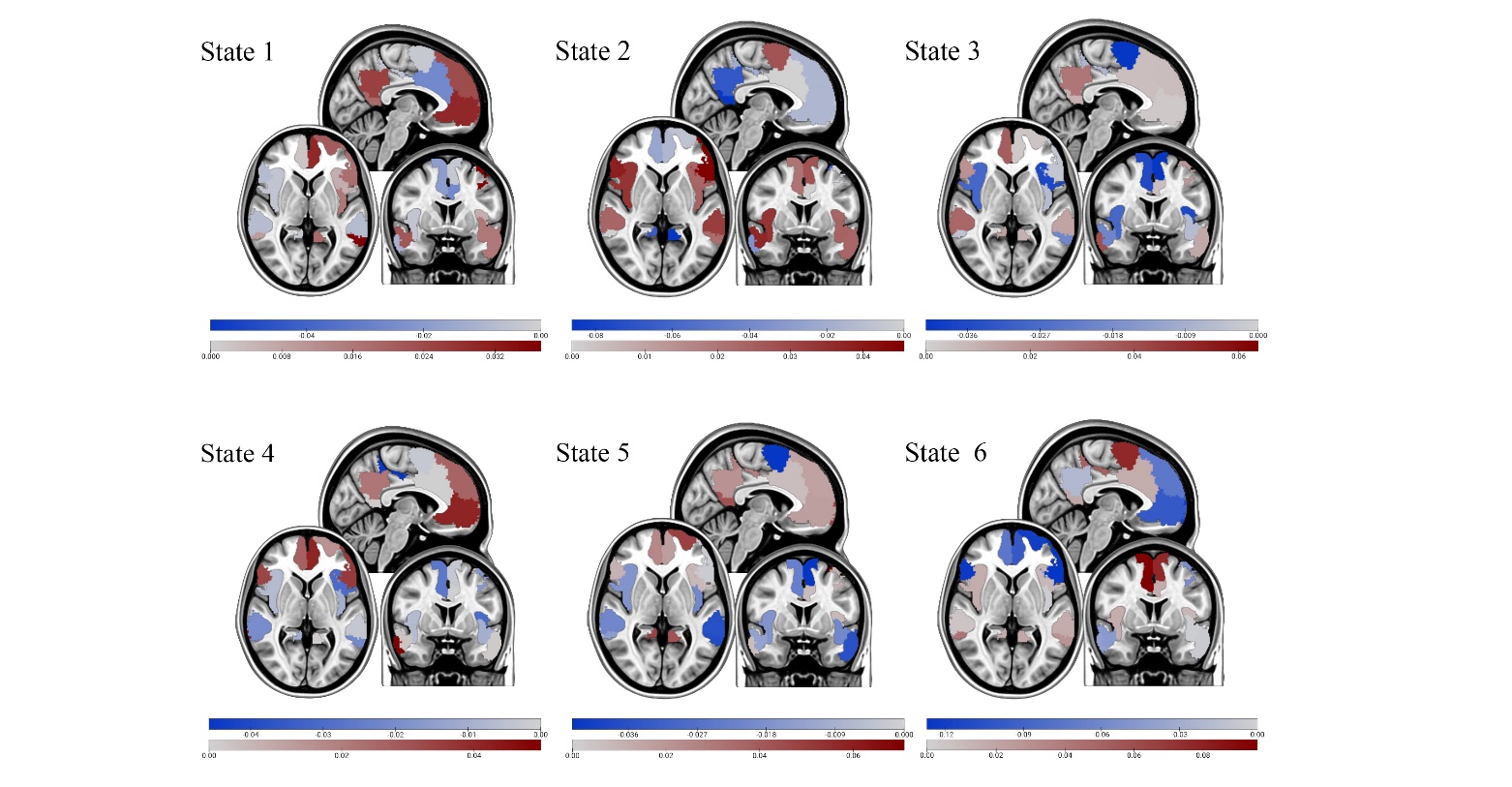

Figure 4. Mean activation maps for each of the six states with the outlier included. Dark blue colors indicate lower activity, grey/white colors indicate normal activity, and dark red colors indicates increased activity. Activity is relative to other states, not to a baseline.

**Mindfulness Meditation Script**

Instructions to be READ: During this session, there will be a focus on using the breath to relax and recover from challenging circumstances. If possible, it is best to breathe through your nostrils. Also, it is possible that you may experience some twitching of muscles and other physical symptoms that seem unusual. These are normal when you relax and are not reason for concern. The practice you are about to experience is very safe.

There are two points I want to emphasize before beginning. First, considerable research has shown that following stressful encounters, people tend to replay in their minds, over and over again, either what they did or did not do. In the following session, rather than focusing on the past, you will be encouraged to notice how you feel right now, to acknowledge these feelings, and then to consciously release tension and to relax your body as best you can. Second, focusing on the breath is a very effective way to calm the body and mind; thus, the breath will emphasized over and over again. So for the next 10-min, simply listen closely to my voice and follow along without passing judgment on whether you are doing what I am asking correctly or not. Let’s begin:

Script: Begin by closing your eyes; making certain that you are lying comfortably in the scanner. Let your hands rest by your side, muscles relaxed, and take 3 or 4 long, slow breaths, feeling your abdomen expand on the inbreath, then notice as it moves toward your spine on the outbreath. Imagine that your abdomen expands like a balloon as the air moves inward; once completing the inbreath, then exhale gradually but completely with little conscious effort on your part.

------------------

20 sec pause

------------------

As you lie here, you may feel impatient or sense that you need to re-position yourself. Rather than doing so, simply make the conscious “choice” to be completely still, to allow your body to be totally supported by the surface beneath you. Notice what it feels like to let go of your body, to be completely relaxed. In fact, you may notice sensations of tingling or warmth, feel a brief muscle spasm, heaviness of a limb, or a feeling of expansiveness. This is simply part of letting go, of relaxing, of being fully connected to whatever is happening moment-to-moment. Feel how the weight of your body is supported by the scanner and let go even more as you pay close attention to each successive outbreath.

------------------

30 sec pause

------------------

With your awareness tuned into the breath, do not be surprised if your body still seems to be tense or your mind remains active with stress-related images, feelings or thoughts. This is quite normal. Notice how stress-related feelings and thoughts are embodied as tension, stiffness, muscle/joint discomfort, and even fatigue—can you make this connection?

What is important is that rather than dwelling on events that have already occurred, suspend all judgment, evaluation, and elaboration of the past; ask yourself: “How does my body feel in reaction to the previous experience?”

The breath is the most natural way to center ourselves, to counter stress, and to relax. It shifts neural activity in the brain to balance the nervous system. So, for the next 30-sec, just lie here quietly and follow the natural rhythm of your breath; pay attention to the body-relate sensations you are feeling right now; releasing them and see if you can create a deeper feeling of relaxation with each outbreath.

------------------

30 sec pause

------------------

At this point, take a moment and scan your body from head to toe, notice points of tension or discomfort with each breath and then release these as you exhale, saying to yourself in your own mind, “Let go.” As you perform this practice of body sanning, notice over and over again what it feels like when you do relax—picking up on any sensation of letting go, of being at-ease, no matter how small or how brief these sensations may last. So to repeat, during the next period of silence, continue to scan your body from head to toe noticing particular points of tension or discomfort and then with each outbreath try to release these are best you can, saying to yourself in your own mind, “let go.”

------------------

20 sec pause

------------------

In the same way, let go of preoccupation and worries, simply being present with each breath, as it moves in and then out. Let your breathing find its own natural rhythm; see if you can notice the cool feeling of the breath as it enters the back of your throat on the inbreath and the warmth as it moves out of your lungs on the outbreath. Sometimes the breath is long or short, sometimes it may be smooth, and other times you may notice irregularities, but whatever you notice just let it be; that is, do not get involved in judging it, mentally commenting on it, or wishing it was different. Simply be open to each breath and accept whatever happens. What sensations related to the breath do you notice during this next period of silent?

------------------

20 sec pause

------------------

No matter how much you focus on your breath, the mind will often kick-up images, feelings or thoughts that can seem disruptive, particularly when we get dragged into replaying the past or worrying about the future. It is unwise to avoid these activities of the mind. Rather, whatever they may be, notice them, try best not to judge or evaluate them, let them pass, and then gently return to awareness of your breath. Again, use the outbreath to release activities of the mind and to recommit yourself to allowing the body to relax completely, refocusing your attention on what it feels like to let go and to be at-ease. For this final period of silence, simply accept whatever activities of the mind you may notice watching each one pass ad you imaging the breath as if was the peaceful ebb and flow of the ocean as it rhythmically moves in and then out.

-------------------

30 sec pause

-------------------

Now that this session is over, take a moment to appreciate the importance of learning how to settle and to relax yourself, particularly following stressful events in daily life. Now open your eyes, move the muscle of your face, and gently contract and relax the muscle of your arms and legs. The technician will be with you shortly.
